# From mice to humans: Cross-species transcriptomics analysis to uncover distinct mechanisms of Diabetic Retinopathy (DR)

**DOI:** 10.1101/2023.10.19.563187

**Authors:** Siddharth Adda, Christophe Legendre, Sampath Rangasamy

## Abstract

Diabetic Retinopathy (DR) is the leading risk for vision loss in working adults worldwide, affecting more than 100 million people. While risk factors such as diabetes duration, hyperglycemia, and dyslipidemia are well-acknowledged, the unifying molecular mechanisms underlying the pathogenesis of DR remain incompletely understood across different species. To bridge this gap, we conducted a comprehensive RNA-sequencing (RNA-seq) meta-analysis on retinal tissue samples from mice, rats, and humans with diabetes. Our integrative bioinformatics analysis elucidated both conserved and species-specific transcriptional landscapes associated with DR. Notably, pro-inflammatory pathways were ubiquitously activated in the retinal tissues across all examined species. However, unique human-specific immune-metabolic signatures emerged, which were not observed in rodent models. In summary, our investigation unveils both universal and species-specific molecular underpinnings of DR, enhancing our understanding of its complex pathobiology. These findings validate the utility of animal models in DR research and underscore the importance of human-focused studies to uncover mechanisms uniquely relevant to human pathology.

## INTRODUCTION

Diabetic Retinopathy (DR) (OMIM# 603933) is the leading cause of blindness in the working-age population ^1^. The onset and progression of DR are often the result of an extended duration of diabetes, lack of glycemic control, increased blood pressure, and dyslipidemia, which lead to an array of retinal lesions causing DR ^2–6^. Animal models for diabetic retinopathy, ranging from the most commonly used rodent models to non-human primate models, are often used to study diabetic retinopathy^3^. Models to study DR are typically induced using drugs (e.g., streptozotocin and alloxan), diets (e.g., high-fat and high-sugar diets), or genetic models (e.g., lepr^db^) to mimic the hyperglycemia and dyslipidemia^3^. Rodents, particularly mice and rats, are the most preferred animal models for investigating the pathogenesis due to cost, short lifespan, rapid onset of DR pathogenesis, and high reproductive rates ^7^. Retinal architecture and vasculature are more complex in humans than in rodents, and the extent of rodent DR pathophysiology is limited ^3,7–9^. Rodent models do not fully recapitulate the vascular proliferative phenotype in the proliferative diabetic retinopathy (PDR) stage ^3,10,11^. Rodents also have smaller retinal surface area and lack a macula but possess greater photoreceptor cell density in the central retina than humans ^12,13.^ In spite, mice and rat serve as valuable models for studying various aspects of diabetic retinopathy (DR) and contributed to the development of FDA-approved clinical treatments for Diabetic Retinopathy (DR). Notably, key molecules such as Vascular Endothelial Growth Factor (VEGF) and Angiopoietin-2 (ANGPT2) have been shown to alter the integrity of the blood-retinal barrier (BRB) in diabetic mice and serve as therapeutic targets for human diabetic retinopathy ^14–16.^

Despite the elucidation of shared pathophysiological and molecular underpinnings of diabetic retinopathy (DR) between humans and model species, a comprehensive understanding remains elusive. Numerous fundamental questions remain unanswered, such as: What conserved molecular mechanisms drive DR across human and model systems? How do these molecular signatures interact across species? What are the distinct species-specific mechanisms that can be utilized to identify novel therapeutic interventions for human DR? Addressing these fundamental questions is essential for advancing our understanding of DR’s underlying mechanisms and developing more effective therapeutic strategies. To answer these questions, and to delineate the utility of model organisms in elucidating the pathogenesis of diabetic retinopathy (DR), we employed an integrated analytical approach to compare the retinal differential gene expression (DEG) of datasets from the NCBI SRA (RNA sequencing data) and GEO dataset (microarray) libraries. Our analysis revealed significant differences in gene expression profiles between the rodent models of DR and human tissue samples, particularly in inflammatory pathways.

## METHODS

To conduct a comprehensive analysis of RNA sequence and microarray data for cross-species analysis, we aggregated raw or processed RNA-seq datasets from multiple species as described below and the detailed flow chart outlined the steps from sample collection, preprocessing, analysis with DESeq2, and downstream functional enrichment analysis (Figure 1).

**Figure 1:**
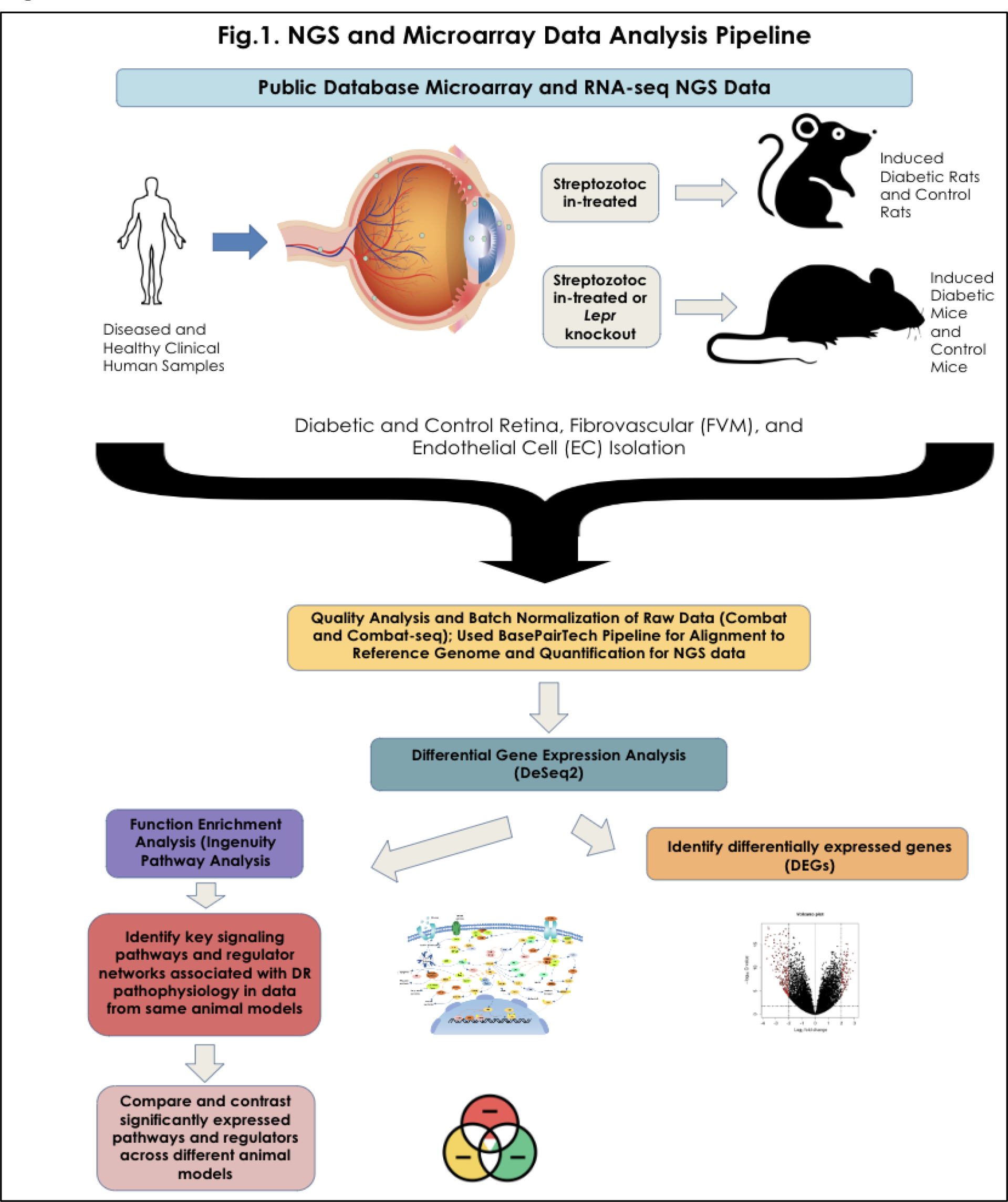
Flow chart of the study’s methodology

### Sample Selection for RNA-sequencing Analysis

Retinal microarray and RNA-seq datasets from mammalian DR models were extracted from the public gene expression databases. Such RNA-seq and microarray datasets were selected/filtered to ensure sample replicate sizes equal to or greater than three, clear descriptions of sample selections, diabetic retinopathy condition (no subtype), sequencing library creation, and processing (separately for each organism type). Samples were ensured to have controlled experimental groups where the diseased condition was the only experimental variable (tissue, age, and animal strain were standardized between the experimental and control groups within a dataset). Other types of RNA-seq studies, such as lncRNAs or miRNAs, were excluded. RNA-seq datasets were also filtered for samples with paired-end reads only. A total of 108 anonymized samples were obtained from the NCBI GEO for microarray datasets (70 datasets) and the SRA for RNA-seq datasets (38 datasets) (post-Principal Component Analysis (PCA) filtering) (Supplemental Table 1). Mice, rats, and human samples of the retina, retinal endothelial cells, and fibrovascular cells with diabetic retinopathy and control conditions were selected. Fastq files from RNA seq were preprocessed using the Base Pairs (basepairtech.com) pipeline, which incorporates quality control using Fastp and read alignment with STAR, which is subsequently used for downstream processing to provide a comprehensive approach for the computational examination of RNA transcript levels.

**Table 1:** Differentially expressed pathways, regulators, and associated gene networks in the analysis of human DR samples.

| Pathway Name | Direction | Top P-value | No. of Genes | Gene Symbols | Description | Enriched Regulator Name | P-value |
| --- | --- | --- | --- | --- | --- | --- | --- |
| Cytokine Storm Signaling Pathway | Upregulated | 1E-19 | 20 | CCL3, CCL4, CCL20, CCL3L1, CCL3L3, COL1A1, COL4A1, COL4A2, COL6A2, CXCL2, CXCL3, CXCL8, FTL, HLA-DRB4, IFNA5, IL10, IRF7, OSM, PPBP, TNFSF13 | Systemic hyper-inflammation caused by high cytokine concentration | TNF | 3.79E-24 |
| S100 Family Signaling Pathway | Upregulated | 4.22E-14 | 15 | ADRA2B, C5AR1, CAMK2B, CCL20, CCL3, CCL4, CCR7, CXCL8, F2R, GPR132, HCAR2, MMP1, NPY4R, NPY4R2, OPN1LW, S100B | Calcium-binding proteins triggering proinflammatory immune response | IFNG | 3.58E-27 |
| Neuroinflammation Signaling Pathway | Upregulated | 2.01E-11 | 12 | CALB1, CCL3, CXCL8, GABRA1, HLA-DRB4, HMOX1, ICAM1, IFNA5, IL10, IRF7, S100B, SLC6A11 | activates proinflammatory cytokines and neurotransmitters | IL6 | 1.97E-19 |
| IL4 Signaling | Upregulated | 5.34E-05 | 9 | COL1A1, COL1A2, COL20A1, COL3A1, COL4A1, COL4A2, COL5A2, COL6A3, COL9A1 | cytokine promoting macrophage and immune activation | VEGF | 9E-09 |
| TREM1 Signaling | Upregulated | 2.22E-08 | 6 | CCL3, CD83, CXCL3, CXCL8, ICAM1, IL10 | wound repair through immunoglobulin receptor with increased expression of chemokines and cytokines | STAT3 | 3.92E-18 |

### RNA-sequencing Analysis

To analyze inter-species variance, datasets with technical replicates were merged and batch-normalized using an empirical Bayesian framework using the Combat and Combat-Seq tools to minimize non-biological variance ^17–19^. Normalization, differential expression analysis, and correction for multiple testing were performed using DESeq2 ^20^. Expression levels were fitted to a negative binomial distribution to calculate the relative transcript expression levels (fold change), normalized between sample groups using the median-of-ratios method, transformed to a log2 scale, and scaled using Bayesian shrinkage estimators (dispersion estimates). The Wald test was used to calculate the P values, which were adjusted using the Benjamini-Hochberg correction to produce adjusted P values (FDR). Subsets of differentially expressed genes (DEGs) in each comparison were selected by filtering out all genes with greater than 0.05 P values and 0.01 FDR adjusted P values (0.1 FDR cutoff for group comparison). The results were validated using a PCA-based quality check and outlier test. Qiagen’s Ingenuity Pathway Analysis (IPA) (QIAGEN Inc., https://digitalinsights.qiagen.com/IPA) ^21^ and Gene Cards ^22^ were used to identify upstream regulators and their effects on cellular pathways.

## RESULTS

To explore the molecular mechanisms differentiating the DR phenotype between rodent models and human cases, we performed an inter-species comparative RNA-seq analysis to identify genes that showed differential expression patterns. We conducted differential gene expression RNA sequencing analysis on 38 RNA sequencing and 70 microarray mouse, rat, and human samples (post-PCA removal) of the retina, retinal endothelial cells, and fibrovascular cells, under diabetic retinopathy and control conditions. The Principal Component Analysis (PCA) quality check to investigate the gene expression variability between replicates, as well as a cluster analysis between experimental groups (Supplemental Figure 1), helped remove approximately 23 samples due to improper clustering. Heatmaps of significant differentially expressed genes in each of the sample comparisons were produced (Supplemental Figure 2-4) Pathway enrichment analysis of differentially expressed genes in human, mouse, and rat samples revealed shared and variably enriched pathways and downstream/upstream regulators between the species. Intra-species sample variation was minimal.

### Differential Gene Expression Analysis of Human Retinal Tissue

Table 1 outlines the pathologically relevant, differentially expressed pathways and regulators between the human retinal samples of diabetes and the control samples. Functional analysis of human samples revealed inflammatory markers of DR, including an upregulated cytokine storm Signaling pathway (p =1E-19), S100 family Signaling pathway (4.22E-14), Neuroinflammation

Signaling pathway (2.01E-11), and Il4 Signaling (5.34E-05). The Cytokine Storm Signaling and Neuroinflammation Signaling Pathways are characterized by a systemic retinal inflammatory response caused by elevated pro-inflammatory cytokine concentrations and the presence of neurotransmitters. The S100 Family Signaling Pathway reflects the upregulation of calcium-binding proteins that activate the inflammatory response alongside interleukin 4 (IL4) signaling, causing a macrophage-centered immune response. Interestingly, we observed a significant downregulation of adipogenic and cholesterol-related pathways (p =1.83E-04) as well as a decrease in the activation pathway of RAR Retinoid (p =1E-02). RAR is often described as an anti-angiogenic factor ^23^ and the reduced activity of RAR has been shown to increase angiogenesis, endothelial cell migration, and reduce adipogenesis. Activation of the Retinoid X Receptor (RXR) ameliorates the progression of diabetic retinopathy in murine models ^24^. Upstream regulators of pro-inflammatory molecular pathways such as TNF-α (p =3.79E-24), VEGF (p =9E-09) that promotes vascular permeability and angiogenesis), immune response-inducing IFNG (p = 3.58E-27), IL6 (p =1.97E-19), and STAT3 (p =3.92E-18) were all significantly upregulated. STAT3, an upstream regulator of a multitude of genes involved in proliferation, differentiation, apoptosis, and inflammation pathways, is also significantly upregulated ^25^.

### Differential Gene Expression Analysis of Mouse Retinal Tissue

Table 2 outlines the pathologically relevant, differentially expressed pathways and regulators between the DR mouse retinal samples of DR compared to the control samples. The analysis of mouse samples revealed a striking difference in pathway expression compared to that in human samples. Both the Cytokine Storm Signaling Pathway (p =1.24E-02) and Wound Healing Pathway (p =7.31E-03) were downregulated. The Wound Healing Inflammatory Pathway regulates and causes angiogenesis and promotes the immune response characteristic of DR and retinal endothelial cell proliferation ^26^. However, the expression of certain pro-inflammatory regulators is shared between mouse and human samples, such as IL1B (p =1.17E-03), TNF (p =1.68E-05), IFNG (p =4.96E-06), and ELAVL1 (p =3.86E-03), which were all upregulated. PPARD (downregulated; p-value:1.99E-02) is involved in lipid metabolism and regulation and insulin secretion, both of which are functions reduced in DR pathophysiology ^27^.

**Table 2:**
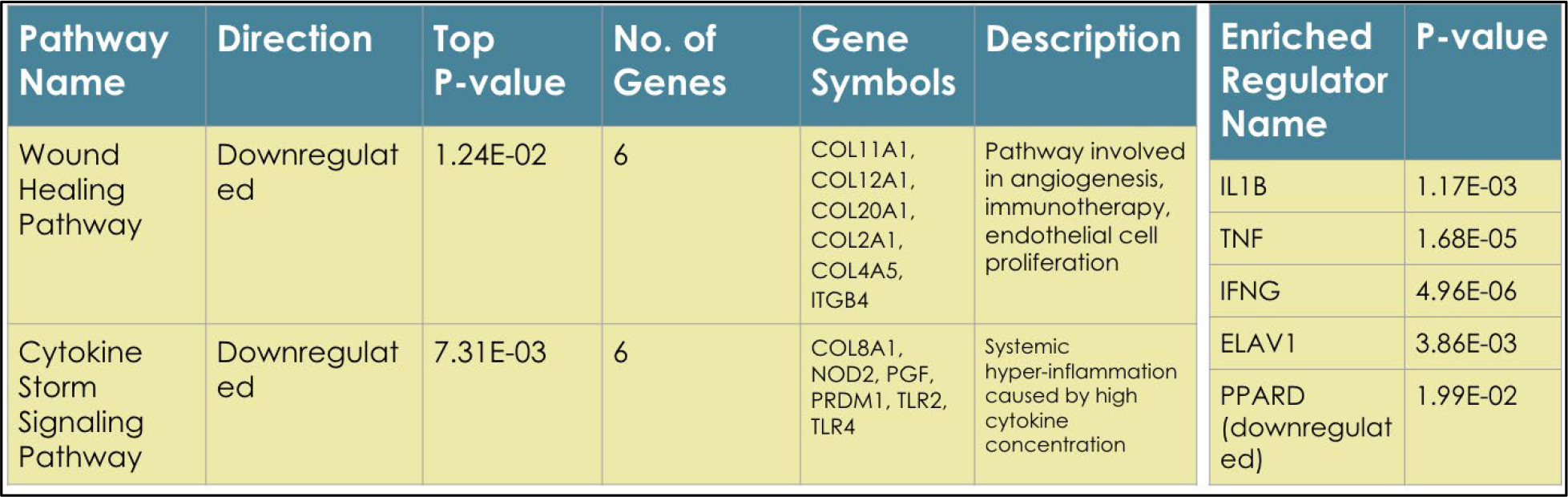
Differentially expressed pathways, regulators, and associated gene networks in the analysis of mice samples.

### Differential Gene Expression Analysis of Rat Retinal Tissue

Table 3 outlines the pathologically relevant, differentially expressed pathways and regulators between the rat retinal samples of DR and control samples. In rat samples, the inflammatory Wound Signaling Pathway (p =2.87E-02), Neuronal CREB Signaling (p =1.91E-02), and Insulin Secretion Pathway (p =2.92E-03) were upregulated. CREB Signaling plays a role in cellular differentiation and glucose homeostasis ^28^. However, the Insulin Secretion pathway was only differentially expressed in the RNA-seq rat samples, not in the microarray samples. Similar to the human samples, the rat samples expressed enriched inflammatory and angiogenic pathways and upstream regulators, IL6 (p-value:2.83E-02) and ATF4 (p-value:2.75E-09), which promote angiogenesis via the regulation of VEGF ^29^, OSM (p-value:4.81E-03), IFNG (p-value:2.86E-02), and FGF2 (p-value:1E-03), which causes vascular dysfunction and angiogenesis ^30^.

**Table 3:**
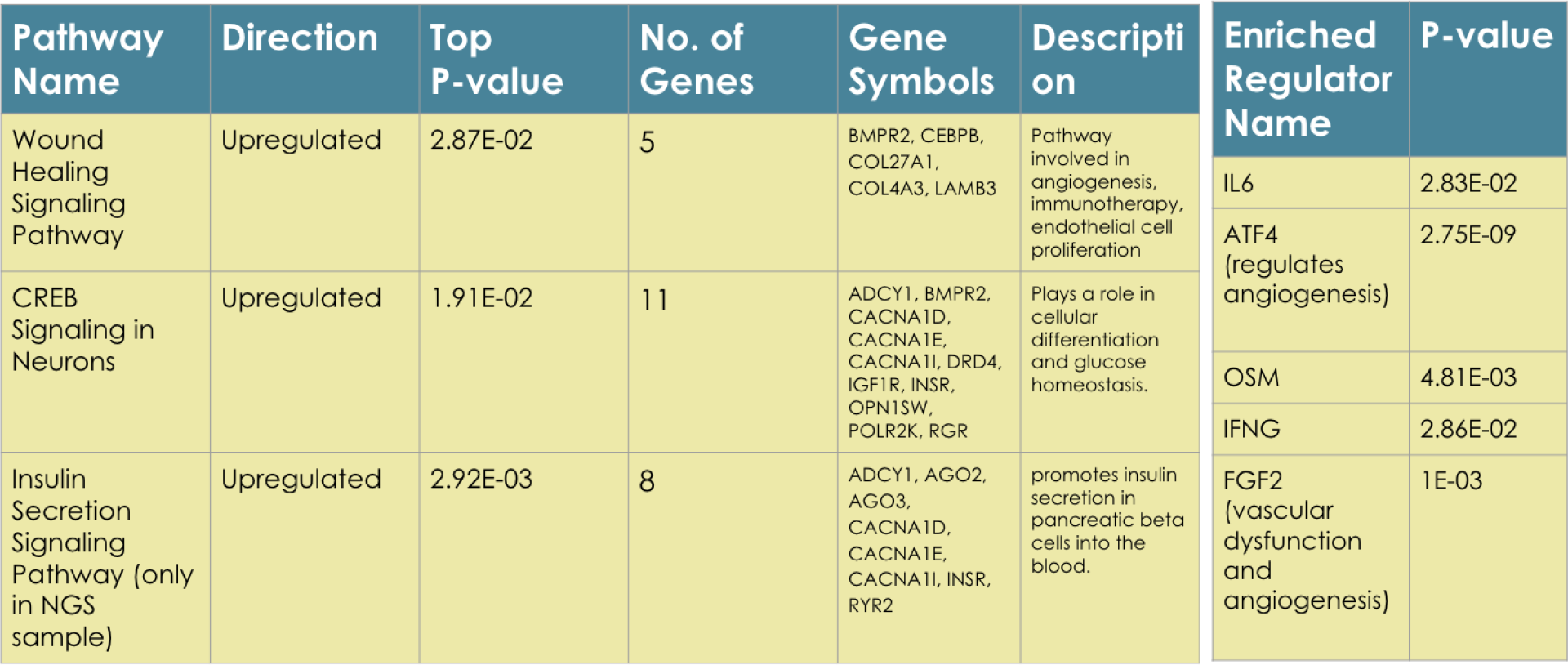
Differentially expressed pathways, regulators, and associated gene networks in the analysis of rat DR samples.

### Cross-Species Functional Analysis

From cross-species comparison analysis, we identified differentially expressed pathways, genes, and regulators in human, mouse, and rat tissue samples. The Venn diagram (Figure 2) describes the overlap in differential expression between rodent models and human samples. The cross-species comparison analysis yielded conserved and variable differentially expressed genes, characteristic of the DR phenotype between the species. Both rat and human samples shared upregulated inflammatory expression signatures (upregulated IL6, cytokine, CREB pathways), but rat expression profiles uniquely had upregulated adipogenic pathways, whereas human samples showed reduced adipogenesis. However, compared with rodent models, human tissue samples have a higher degree of expression and a greater variety of factors related to inflammation. Mouse models showed comparatively reduced inflammation, but still shared key upregulated molecular markers of DR in mice, rats, and human samples, such as TNF-α, IFNG, and VEGF.

**Figure 2:**
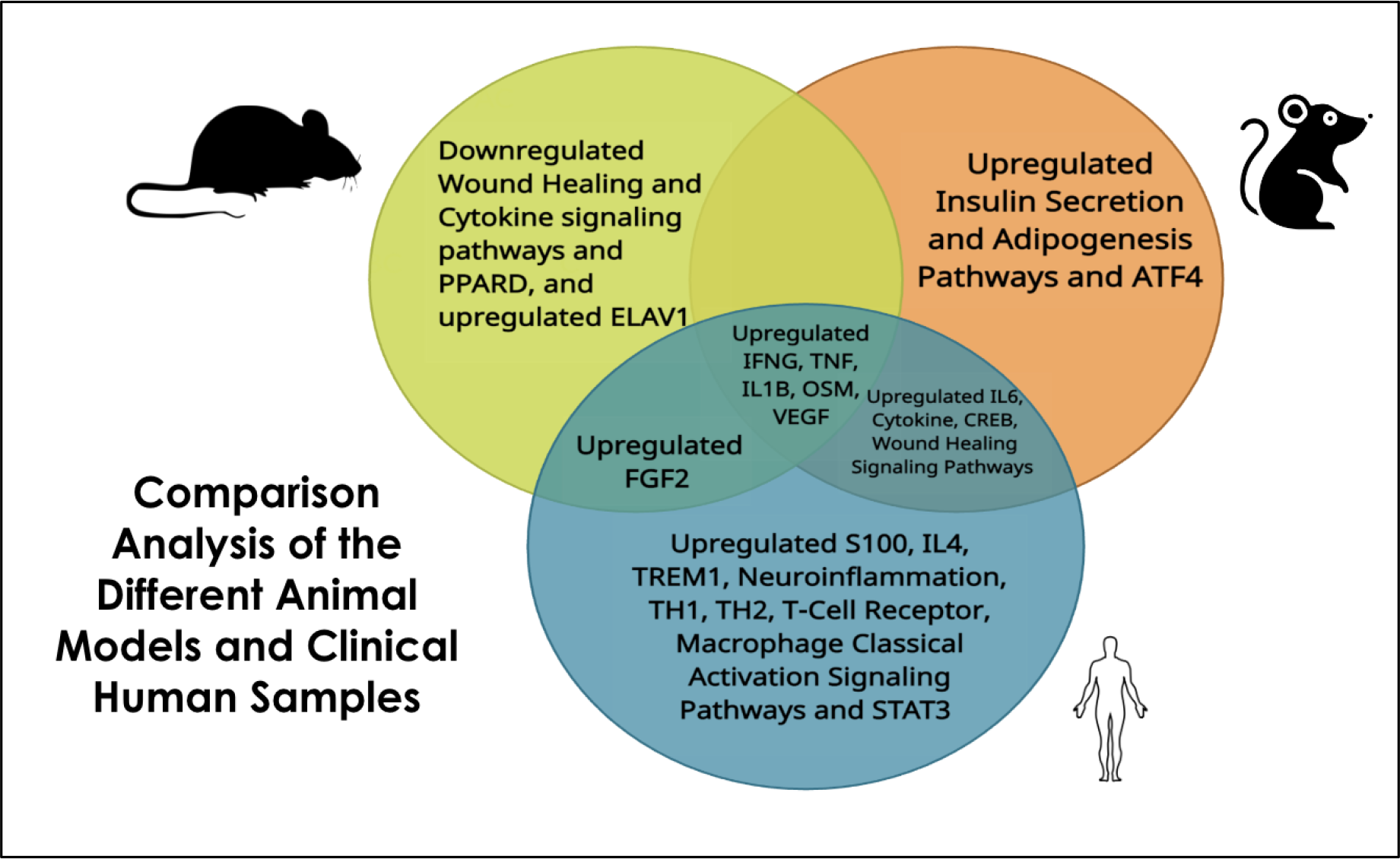
Venn diagram comparing pathologically significant enriched pathways and regulators between rodent models of DR and human DR.

## DISCUSSION

The study provides strong evidence of the significant differences in retinal gene expression profiles between rodent models and human diabetic retinopathy. The rat models show an excessive expression of insulin secretion and adipogenic pathways, while mouse models have a unique downregulation of certain pro-inflammatory pathways. Despite these differences, several crucial angiogenic and pro-inflammatory cytokine pathways display a high degree of conservation across species. Interestingly, the human DR model displays a distinct pathophysiological landscape, characterized by heightened angiogenesis and inflammation, compared to its rodent counterparts. Notably, certain cytokines, such as IL1B, and the angiogenic factor VEGF, are ubiquitously expressed but exhibit variations in upregulation intensity.

Rodent models are commonly used to understand the underlying molecular mechanisms and speed up the development of treatments for diabetic retinopathy^3^. Although these models are well-established and cost-effective for identifying and evaluating DR therapies, their usefulness in human research is still unclear. While rodent models can capture some molecular characteristics of human DR, they cannot recapitulate the full spectrum of human DR pathophysiology as our analysis indicate. This is because numerous factors contribute to the phenomenon, with one of the main factors being the inability of these models to accurately mimic the pathophysiological complexities associated with both type 1 and type 2 diabetes^31^. Our observations also highlights that the mouse models have lower expression of inflammatory pathways, which is because mice have been found to have greater resistance to inflammatory challenges such as DR than humans^32^. However, this study also showed that rat models of DR recapitulate some of the inflammatory phenotypes exhibited by human samples. Multiple factors collectively influence the gene expression pattern across diverse species in a tissue. Notably, the unique composition, disease complexity, genetic homogeneity, and evolutionary divergence in immune systems between mice, rats, and humans may contribute to distinct gene expression variations in the model systems.

Our study highlights the paucity of readily available RNA-seq and microarray datasets that meet stringent criteria for sample descriptions, controlled experimental designs devoid of extraneous treatment conditions, and explicit identification of mice/rat species types. While we employed a stringent selection process to identify datasets with high similarity, we encountered significant heterogeneity, particularly in mouse datasets, attributable to strain differences, variability in diabetic induction methods, disparities in diabetes duration, and inconsistencies in sex descriptions. The inclusion of multiple strains of both mouse and rat samples in our study (C57BL/6 streptozotocin-induced mice, Lepr^-/-^, Ins2 Akita, Sprague Dawley and Long Evans streptozotocin-induced rats) further underscores the inherent variability associated with rodent models of diabetic retinopathy. In contrast, human datasets introduced additional variables for consideration, such as the inclusion of fibrovascular and endothelial cells in addition to retinal data, as well as a more limited sample size compared to rodent models. To mitigate the potential for publication bias, we opted to include all datasets in our analysis, thereby enhancing its comprehensiveness and reliability. Furthermore, we placed a strong emphasis on filtering low-quality RNA-seq data using a rigorous strategy, as previously advocated.

In summary, our findings indicate notable similarities and alsovariations in diabetic retinas of mice, rats, and humans. This study revealed that a significant proportion of the genes that exhibited overexpression were associated with the inflammatory response, which is a distinguishing feature of diabetes in rodent models of DR. The animal models and clinical samples showed minimal variance within species, but there was variation observed between animal models across different species. Based on both rat models and human samples, there was a significant overlap in the inflammatory and angiogenic pathways. However, in mouse models, there appears to be a disparity in the expression of these inflammatory pathways.

Further research and acquisition of additional data are necessary to substantiate the observed physiological differences within and between species, which are crucial in identifying the targets and biomarkers relevant to DR development.

## Ethical Compliance

Analysis performed in studies did not involve any human participants.

## Data deposition and access

Data Access Statement: Research data supporting this publication are available in NCBI SRA and GEO dataset repositories according to Supplement Table 1. Unpublished data used in the study will be updated in appropriate data repositories in the near future.

## Acknowledgments

Supported by the TGen-Sylvia Chase Early Career Training Program award.

## Author Contributions

SA and SR contributed to the design and implementation of the research, CL helped in the sequencing process and all authors contributed to the writing of the manuscript. SR conceived the original idea and supervised the project.

## Notes

In this article, we have represented human, mouse, and rat genes based on the HUGO Gene Nomenclature Committee (HGNC) standard (Gene name all in capital letter), rather than using species-specific gene nomenclature. We believe this approach ensures clarity and consistency in cross-species analysis when referring to gene names.

