## Supplement Table for "From mice to humans: Cross-species transcriptomics analysis to uncover distinct mechanisms of Diabetic Retinopathy (DR)"

### SUPPLEMENTS

**Supplement Figure 1:** Three PCA graphs of human, mouse, and rat samples.

**Supplement Figure 2:** Heatmaps of differentially expressed genes in human samples and multiple comparisons.

**Supplement Figure 3:** Heatmaps of differentially expressed genes in mouse samples and multiple comparisons.

**Supplement Figure 4:** Heatmaps of differentially expressed genes for rat samples.

**Supplement Table 1:** RNA-seq and microarray samples used in this study.

**References**

**Supplement Figure 1**

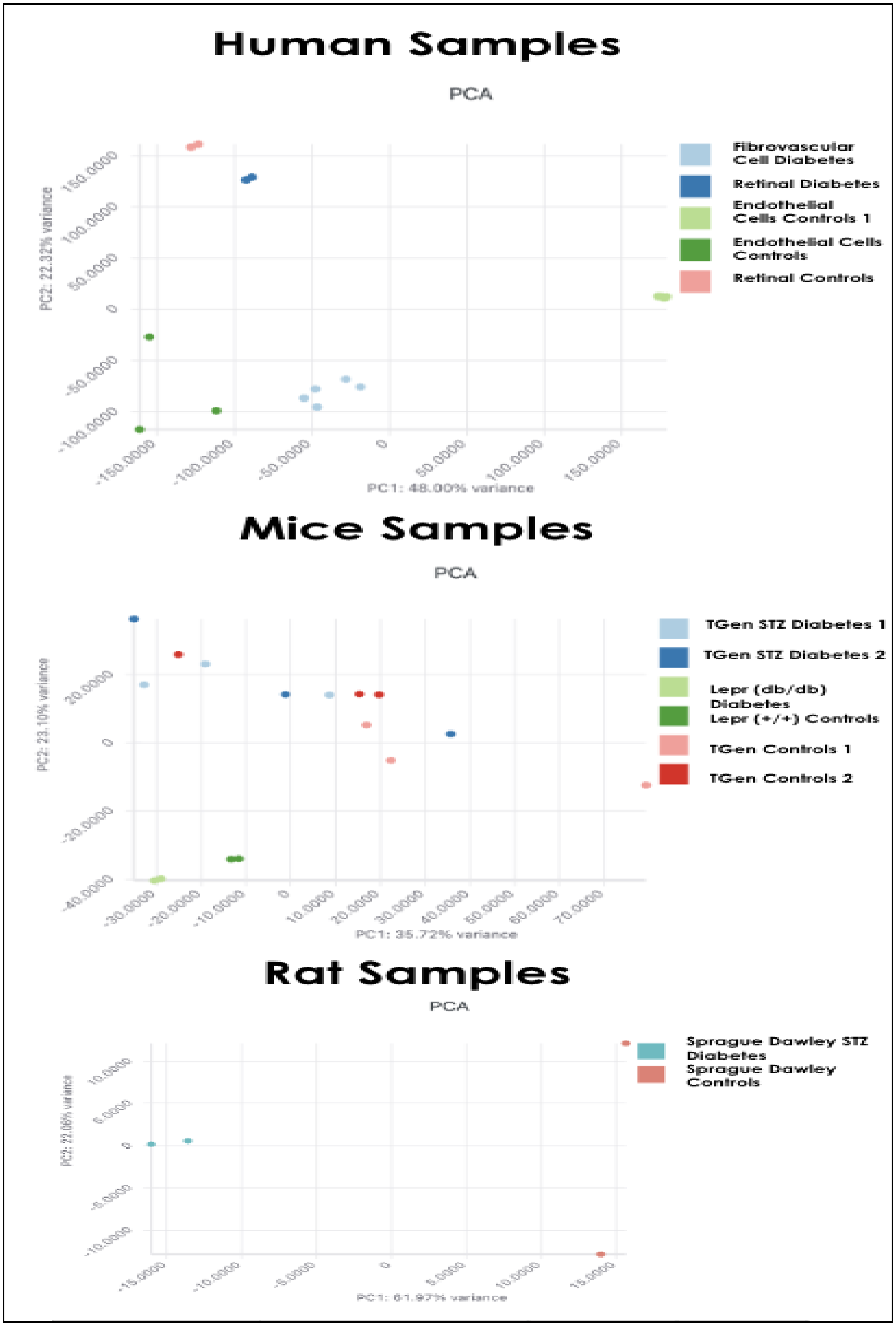

**Supplement Figure 2**

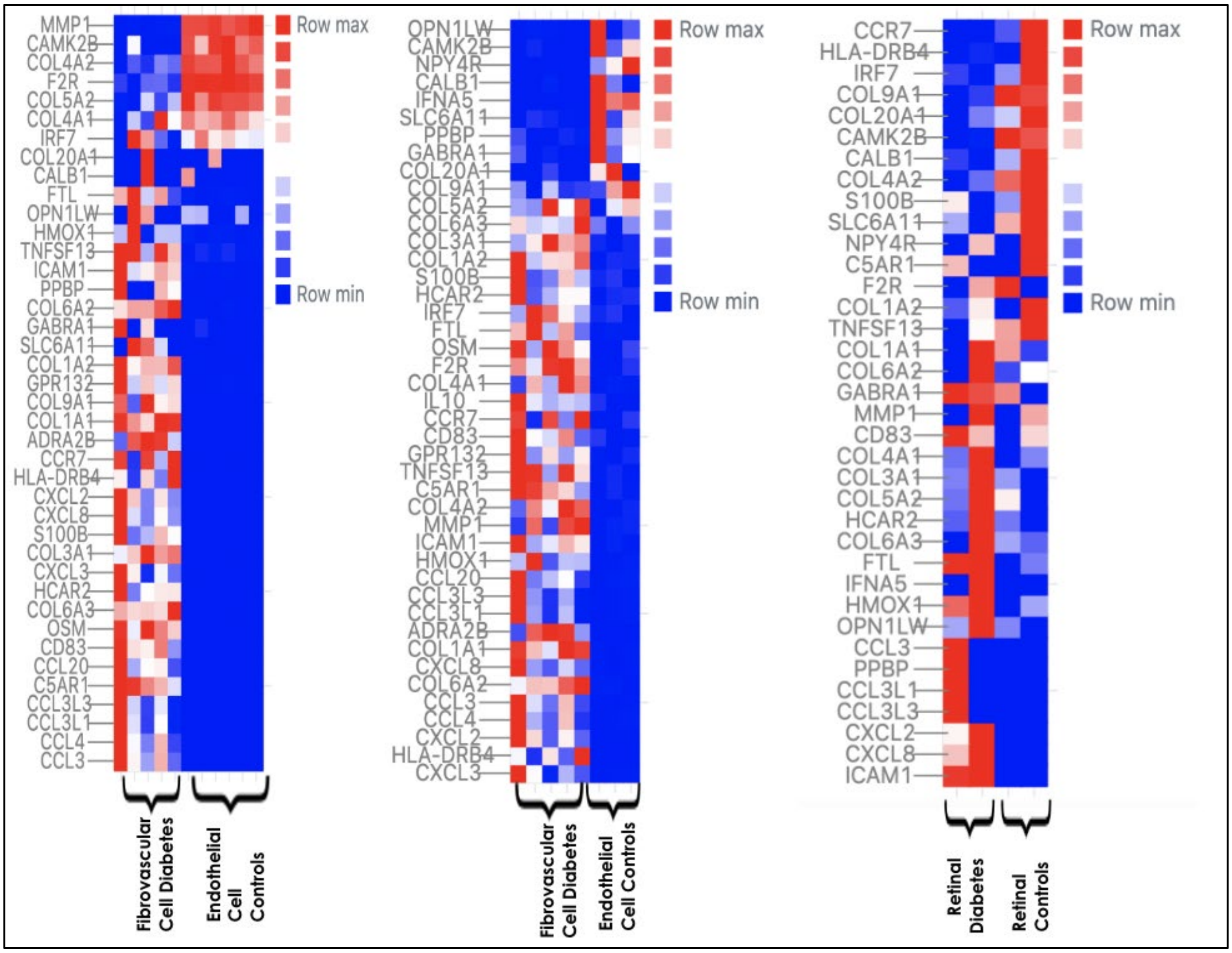

**Supplement Figure 3:**

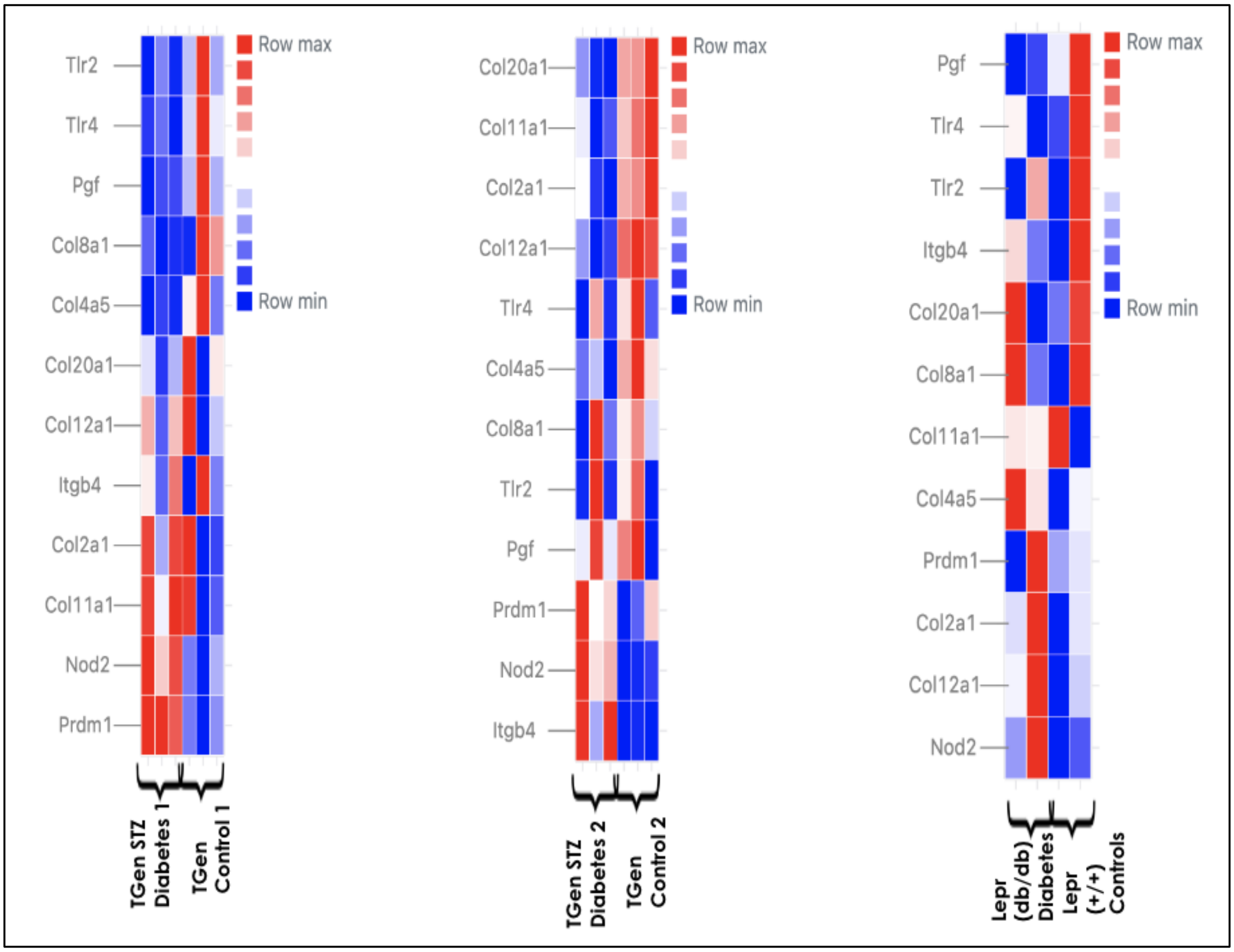

**Supplement Figure 4:**

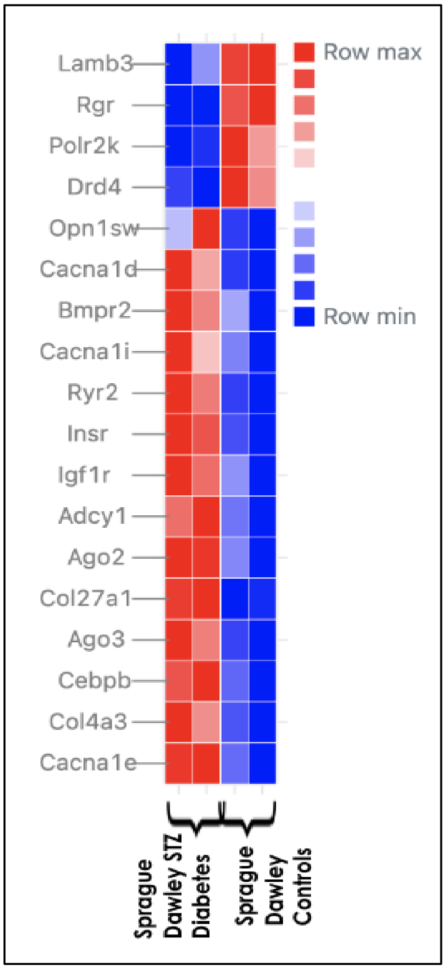

**Supplement Table 1:**

| Name | SRA/GEO Ids | Sample/Tissue Type | Data Type | No. of Samples |
|---|---|---|---|---|
| Lam et al. 2017 | SRP097696 | Human FVM Diabetes & Endothelial Cell Controls | RNA-seq | 8 |
| Huang et al. 2019 | SRP239417 | Human Retinal Endothelial Cell Controls | RNA-seq | 6 |
| Li et al. 2019; Luo et al. 2022 | SRP115195 | Human Retina Diabetes and Control | RNA-seq | 4 |
| Unnamed | SRP323188 | Sprague Dawley Rat Diabetes and Controls | RNA-seq | 4 |
| Chen et al. 2021 | SRP281889 | Lepr Knockout Mice Diabetes and Controls | RNA-seq | 4 |
| Unpublished | NA | Mice Diabetes and Controls | RNA-seq | 12 |
| Friedrichs et al. 2017 | GSE87433 | C57BL/6J Mice Diabetes and Controls | Microarray | 12 |
| Freeman et al. 2009 | GSE19122 | INS2 AKITA/+ vs C57BL/6G mice Diabetes and Controls | Microarray | 25 |
| Kirwin et al. 2011 | GSE28831 | Long Evans Rat Diabetes and Controls | Microarray | 18 |
| Bixler et al. 2011 | GSE24423 | Sprague Dawley Rat Diabetes and Controls | Microarray | 9 |
| Ishikawa et al. 2015 | GSE60436 | Human FVM Diabetes and Controls | Microarray | 6 |
